# Expression patterns of PD-L1 and PD-1 provide rationales for immune checkpoint inhibition in soft tissue sarcomas

**DOI:** 10.1101/569418

**Authors:** Martin F. Orth, Veit L. Buecklein, Eric Kampmann, Marion Subklewe, Elfriede Noessner, Florencia Cidre-Aranaz, Laura Romero-Pérez, Fabienne S. Wehweck, Lars Lindner, Rolf Issels, Thomas Kirchner, Annelore Altendorf-Hofmann, Thomas G. P. Grünewald, Thomas Knösel

## Abstract

Soft tissue sarcomas (STS) are highly malignant cancers with mesenchymal origin. In many instances, clinical outcome is poor due to high rates of local recurrence and metastasis.

The programmed death receptor ligand 1 (PD-L1) is expressed in several cancers. PD-L1 interacts with its receptor, PD-1, on the surface of tumor infiltrating lymphocytes (TILs), thereby attenuating anti-cancer immune response. Immune checkpoint inhibitors targeting this interaction are promising new anti-cancer drugs. However, present studies on the PD-L1 and PD-1 expression status in STS are limited either by small sample size, analysis of single STS subtypes, or lack of combinatorial assessment of PD-L1, PD-1 and TILs.

To overcome these limitations, we evaluated the expression patterns of intratumoral PD-L1, the amount of TILs and their PD-1 expression status, as well as associations with clinicopathological parameters in a large and comprehensive cohort of 274 samples comprising more than six STS subtypes.

We found that nearly all STS subtypes showed partial PD-L1 expression, albeit with a broad range of PD-L1 positivity across subtypes (50% angiosarcomas, 23% UPS, 13% leiomyosarcomas, 12% dedifferentiated liposarcomas, 3% synovial sarcomas, 0 MPNST, and 18% mixed sarcomas). Co-expression and correlation analyses uncovered that expression of PD-L1 was associated with more PD-1 positive TILs (*P* < 0.001), higher tumor grading (*P* = 0.022) and worse patients’ 5-year overall survival (*P* = 0.016).

In sum, the substantial portion of STS showing PD-L1 expression, the simultaneous presence of PD-1 positive TILs, and the association of PD-L1 with unfavorable clinical outcome provide a rationale for immune checkpoint inhibition in patients with PD-L1-positive STS.

## INTRODUCTION

Soft tissue sarcomas (STS) are heterogeneous and highly malignant tumors originating from the mesenchymal lineage^1^ with more than 50 subtypes described to date^2^. Current therapy regimens for STS are limited mainly to surgery and radiation^3^. Benefits of neoadjuvant and adjuvant radiochemotherapy are still under debate^3,4^. In addition, established therapies appear not fully sufficient for long-time tumor control as many patients develop local relapse (up to 45%) and/or distant metastases (30%)^1,5,6^, leading to fatal outcome.

Unfortunately, STS patients barely benefited from new and more sophisticated anti-cancer treatments like kinase inhibitors^3^, and innovative and more effective therapeutic alternatives are lacking, possibly due to the rarity and diversity of STS.

In the past two decades, immune checkpoint inhibitors revolutionized anti-cancer therapies. Immune checkpoint inhibitors like Nivolumab interfere with an immunosuppressive mechanism by which cancer cells attenuate the anti-cancer activity of the patient’s immune system^7^. Specifically, several cancer cells hijack a regulatory mechanism of the immune system by expression of programmed death receptor ligand 1 (PD-L1, B7-H1, CD274)^8^. PD-L1 is normally expressed on antigen presenting cells and binds to the programmed death receptor 1 (PD-1) on activated T cells, B cells, and macrophages, thereby blocking their activity and the recruitment of further immune cells^9^. In melanoma and urothelial carcinoma with PD-L1 expression, inhibition of this immune checkpoint enhances anti-tumor immune activity resulting in significantly improved clinical outcome^10,11^.

However, to date there is not any extensive study on STS that has comprehensively investigated whether PD-L1 expression is a common feature in STS, if PD-L1-positive STS exhibit tumor infiltrating lymphocytes (TILs), and whether those TILs are positive for PD-1 as a prerequisite of PD-L1 and PD-1 interaction^9,12–14^.

In the present study, we analyzed these aspects in a large tissue microarray (TMA) comprising 225 STS samples of six distinct subtypes and 22 additional samples of various other STS subtypes. Our results show that a substantial proportion of STS is positive for PD-L1 and that PD-L1 expression is associated with PD-1 positive TILs, and poor patient outcome, providing a rationale for immune checkpoint inhibition in these STS patients.

## MATERIALS AND METHODS

### Patient cohort

Formalin fixed paraffin-embedded tumor material from STS patients was retrieved from the archive of the Institute of Pathology, LMU Munich (years 1989 to 2012) in agreement with the ethics committee of the LMU Munich University hospital. Tumors were reclassified by TKn and EK according to the current WHO-classification. Clinical data, including sex, age at diagnosis, tumor site, tumor size, metastasis, and grading were extracted from the archived pathological results and the database of the department for hyperthermia treatment of the LMU Munich. Survival data were updated until 09/2017 in collaboration with clinicians performing follow-up and the patients’ respective general physicians. The resulting cohort comprised 247 STS cases (**Table 1**).

**Table 1:** Patients’ characteristics

| **STS subtype** | ***n*** | **Fraction of cohort [%]** | **Sex (male / female)** | **Age at diagnosis (median, range) [years]** | **Follow-up (median, range) [months]** |
| --- | --- | --- | --- | --- | --- |
| Angiosarcoma | 6 | 2.4 | 3 / 3 | 48 (18-67) | 14 (4-54) |
| Leiomyosarcoma | 47 | 19.0 | 17 / 30 | 55 (19-79) | 33 (6-287) |
| Dediff. Liposarcoma | 49 | 19.8 | 30 / 19 | 57 (24-79) | 65 (0-189) |
| MPNST | 11 | 4.5 | 7 / 4 | 37 (24-74) | 20 (6-102) |
| Synovial sarcoma | 29 | 11.7 | 11 / 18 | 42 (22-67) | 54 (2-176) |
| UPS | 83 | 33.6 | 56 / 37 | 56 (19-79) | 32 (1-182) |
| Mixed | 22 | 8.9 | 9 / 13 | 50 (18-72) | 24 (2-165) |
| **All samples** | **247** | **100.0** | **123 / 124** | **53 (18-79)** | **36 (0-287)** |

### Assembly of TMAs

For TMA assembly, representative tumor areas were marked on hematoxylin and eosin (H&E) stained slides of the selected paraffin blocks of each patient, and two cores with 0.6 mm diameter were taken from each sample. Samples from tonsils were added as positive controls.

### Immunohistochemistry and scoring of immunoreactivity

TMA sections of 5 μm were stained for H&E, PD-L1, PD-1, and Ki-67. Immunohistochemical staining for PD-L1, PD-1, and Ki-67 was performed automatically on a Ventana Benchmark XT autostainer system with the XT ultra-View DAB Kit (Ventana Medical Systems, Roche, Basel, Switzerland). Primary antibodies were the monoclonal anti-PD-L1 antibody raised in rabbit (1:100; E1L3N, Cell Signaling Technology, Danvers, MA, USA), monoclonal mouse anti-PD-1 antibody (1:80; 315M-96, MEDAC, Wedel, Germany), and monoclonal mouse anti-Ki-67 antibody (M7240, Agilent, Santa Clara, CA, USA). Hematoxylin (Vector Laboratories, Burlingame, CA, USA) was used for counterstaining.

PD-L1 expression was scored semi-quantitatively by a pathologist (TKn) and a physician experienced in immunohistology (EK) independently with the categories no (0), weak (1), intermediate (2), and strong (3) immunoreactivity. Only if >1% tumor cells exhibited membranous staining for PD-L1, the corresponding sample was considered as PD-L1 positive. TILs were count per high power field (HPF) (400x magnification) in H&E stained TMA slides. PD-1 positive TILs were quantified in the same way using the PD-1 stained TMA slides. In case of discrepancies in the scoring results of both investigators, consent was built after individual reevaluation of each sample. The percentage of Ki-67 positive cancer cells was evaluated by three researchers (FCA, FW, LRP) independently, and the average percentage of Ki-67 positivity for each sample was taken as basis for further analysis. All researchers scoring the TMAs were blinded to the clinical data.

### Statistical analyses

For statistical analyses PD-L1 expression was classified as negative (0) or positive (>1% of tumor cells with staining intensity 1-3). Samples with ≥4 TILs per HPF were considered as positive for TILs, and if ≥4 TILs exhibited PD1 staining, as PD-1 positive. Statistical analyses were carried out and displayed using SPSS and GraphPad PRISM (v5). Associations between clinicopathological parameters and histological results were calculated with the Fisher’s exact test and unpaired two-tailed Student’s *t*-test. Associations with survival were displayed with the Kaplan-Meier method and significance was assessed with the log-rank test. *P* values < 0.05 were considered statistically significant.

## RESULTS

### PD-L1 is expressed in several STS subtypes

To test whether PD-L1 expression at protein level is a common feature of STS, we stained TMAs with 247 STS samples represented as duplicates by immunohistochemistry (**Table 1**) using an automated immunostainer and a well-established routine antibody for PD-L1, and considered all samples with >1% membranous staining as PD-L1 positive, the remaining as negative (**Figure 1**). The fraction of positive samples comprised 50% of angiosarcomas, 23% of undifferentiated pleomorphic sarcomas (UPS), 13% of leiomyosarcomas, 12% of dedifferentiated liposarcomas, 4% of synovial sarcomas, 0 MPNST, and 22% of mixed sarcomas (**Table 2**). On average, 16% of the entire cohort exhibited PD-L1 expression.

**Figure 1:**
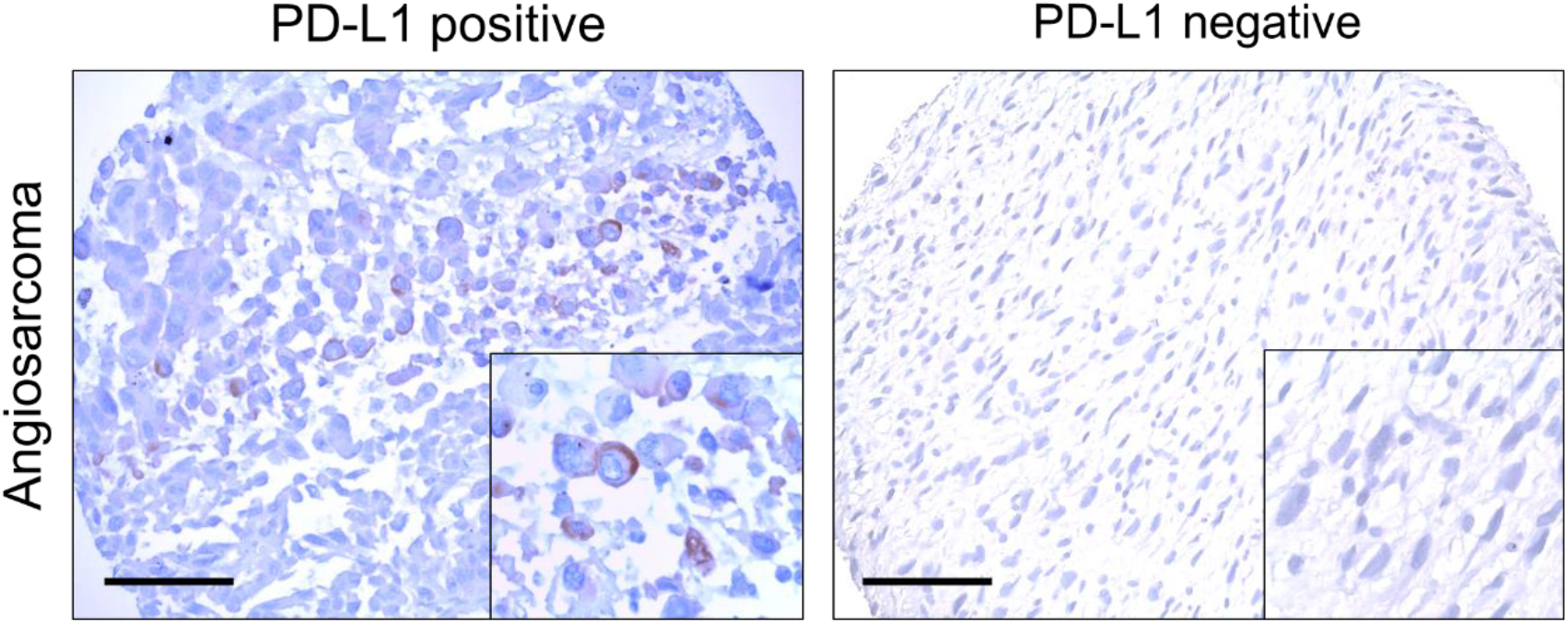
PD-L1 is expressed in a fraction of STS. Representative micrographs of cores on a TMA representing angiosarcoma samples, immunohistochemically stained for PD-L1 (brown). Scale bar indicates 100 μm.

**Table 2:**
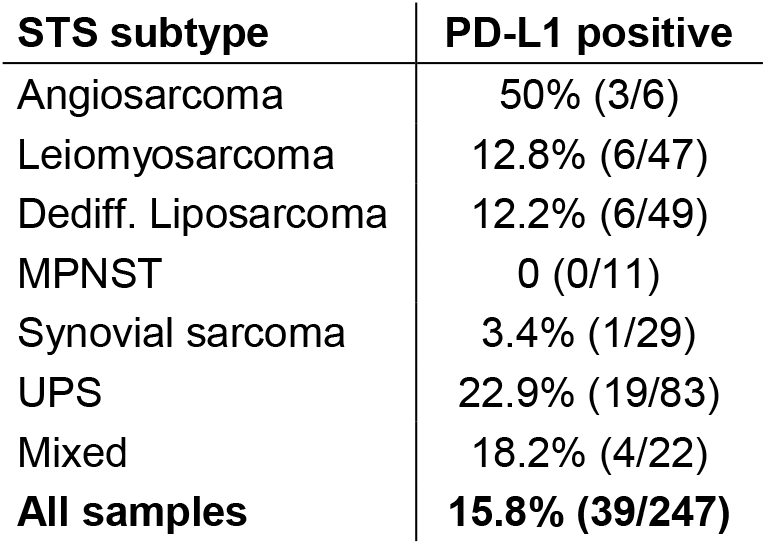
PD-L1 expression in STS

| **STS subtype** | **PD-L1 positive** |
| --- | --- |
| Angiosarcoma | 50% (3/6) |
| Leiomyosarcoma | 12.8% (6/47) |
| Dediff. Liposarcoma | 12.2% (6/49) |
| MPNST | 0 (0/11) |
| Synovial sarcoma | 3.4% (1/29) |
| UPS | 22.9% (19/83) |
| Mixed | 18.2% (4/22) |
| **All samples** | **15.8% (39/247)** |

Hence, a substantial fraction of STS tumors expresses PD-L1 at protein level, albeit with variable proportions depending on the STS subtype.

### PD-L1 expression in STS correlates with PD-1 positive TILs

Besides PD-L1 expression, another prerequisite for effective immune checkpoint inhibition in cancer is an actual interaction of PD-L1 with its receptor. Accordingly, we scored the total number of TILs and TILs positive for PD-1 for all samples tested for PD-L1 positivity considering samples with counts of ≥4 lymphocytes per HPF as positive (**Figure 2**).

**Figure 2:**
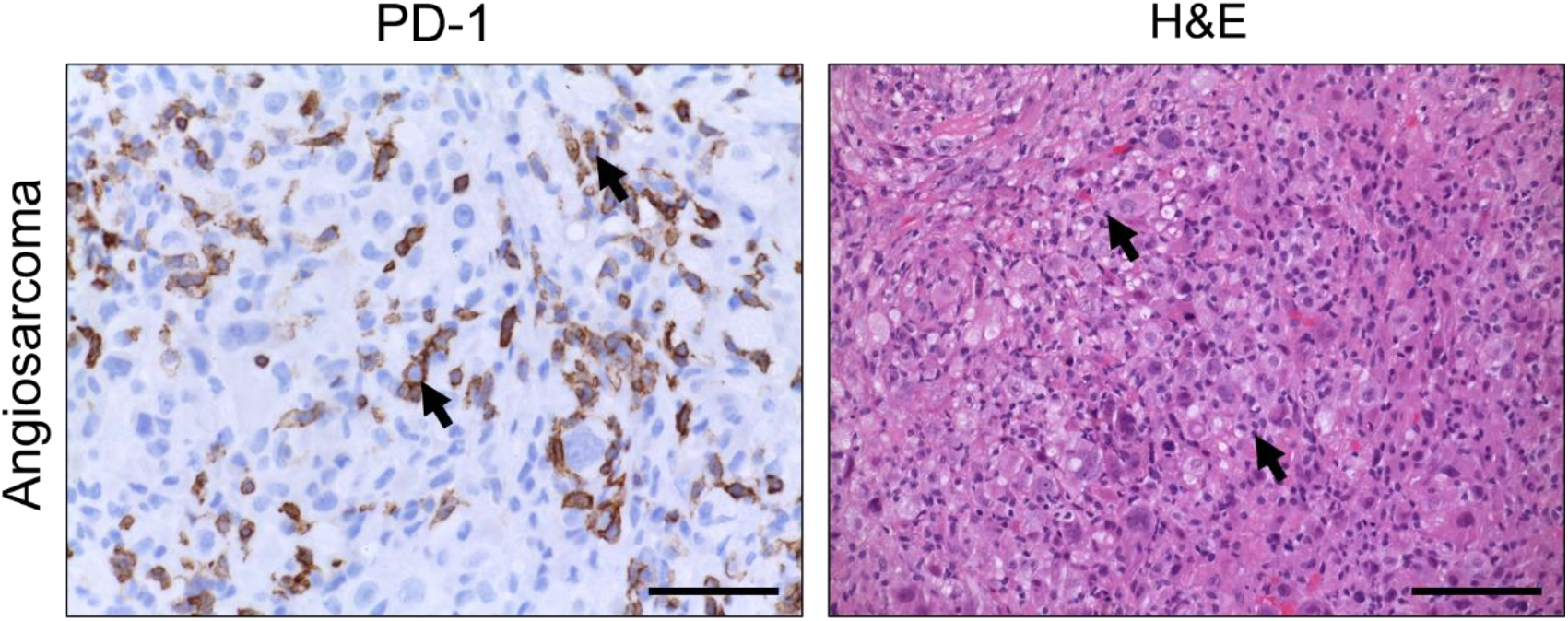
PD-L1 expressing STS are often positive for PD-1 and TILs. Representative micrographs of cores on a TMA representing angiosarcoma, immunohistochemically stained for PD-1 (brown) and H&E. Scale bar indicates 50 μm, arrow-heads point to lymphocytes.

From all 247 samples, sufficient material was available for 245 and 242 samples to evaluate the number of TILs and their PD-1 immunoreactivity, respectively. Across the entire cohort 76.3% of samples were positive for TILs and 28.1% for PD-1. Interestingly, in the PD-L1 positive samples the fractions for TIL and PD-1 positivity were 87.2% and 62.9% (**Table 3**). In fact, for all tested STS subtypes with ≥6 PD-L1 positive samples, the fraction of PD-1 positivity was higher in the PD-L1 expressing samples than in those negative for PD-L1. Consistently, PD-L1 expression and PD-1 positivity were highly significantly correlated (*P* < 0.001, **Table 3**).

**Table 3:** PD-1 positive cells and TILs in STS

| STS subtype | PD-1 positive | PD-1 in PD-L1 positive | P value association PD-L1 - PD-1 | TIL positive | TIL in PD-L1 positive | P value association PD-L1 - TILs |
| --- | --- | --- | --- | --- | --- | --- |
| Angiosarcoma | 40% (2/5) | 33.3% (1/3) | 1.000 (FE) | 83.3% (5/6) | 66.7% (2/3) | 1.000 (FE) |
| Leiomyosarcoma | 17.4% (8/46) | 66.7% (4/6) | 0.006 (FE) | 65.2% (30/46) | 66.7% (4/6) | 1.000 (FE) |
| Dediff. Liposarcoma | 18.8% (9/48) | 66.7% (4/6) | 0.008 (FE) | 81.3% (39/48) | 100% (6/6) | 0.578 (FE) |
| MPNST | 27.3% (3/11) | NA | NA | 81.8% (9/11) | NA | NA |
| Synovial sarcoma | 10.3% (3/29) | 0% (0/1) | 1.000 (FE) | 48.3% (14/29) | 0% (0/1) | 1.000 (FE) |
| UPS | 46.9% (38/81) | 61.1% (11/18) | 0.192 (FE) | 85.5% (71/83) | 94.7% (18/19) | 0.280 (FE) |
| Mixed | 22.7% (5/22) | 50% (2/4) | 0.210 (FE) | 86.4% (19/22) | 100% (4/4) | 1.000 (FE) |
| **All samples** | **28.1% (68/242)** | **62.9% (22/35)** | **<0.001 (FE)** | **76.3% (187/245)** | **87.2% (34/39)** | **0.101 (FE)** |

FE: Fisher's exact test.

Taken together, these results provide evidence that PD-L1 positive STS are enriched for PD-1 positivity, pointing to an actual interaction of PD-L1 and PD-1 positive TILs in STS, which is a prerequisite for patients’ eligibility for immune checkpoint inhibitor therapy.

### PD-L1 expression is associated with clinical outcome in STS

To test whether the PD-L1 status is associated with clinicopathological parameters, we correlated the PD-L1 scoring results obtained from assessors blinded to the clinical data with the most important prognostic parameters for STS patients.

While PD-L1 expression did not correlate with age and tumor size in any tested STS subtype, it was significantly enriched in males (*P* = 0.009) and significantly associated with the prognostically favorable tumor localization in the extremities (*P* = 0.006). In contrast, PD-L1 expression was significantly associated with the prognostically unfavorable parameters of high grading (*P* = 0.022), metastasis at diagnosis (*P* = 0.021), and higher rates of proliferating cells (assessed by percentage of Ki-67 positive tumor cells, *P* = 0.046) (**Table 4**). In accordance, intratumoral PD-L1 expression was associated with a significantly worse 5-year overall survival (37% vs. 58% 5-year overall survival probability, *P* = 0.016) (**Table 4, Figure 3**). In synopsis, these data indicate that PD-L1 expression is clinically relevant in STS patients.

**Figure 3:**
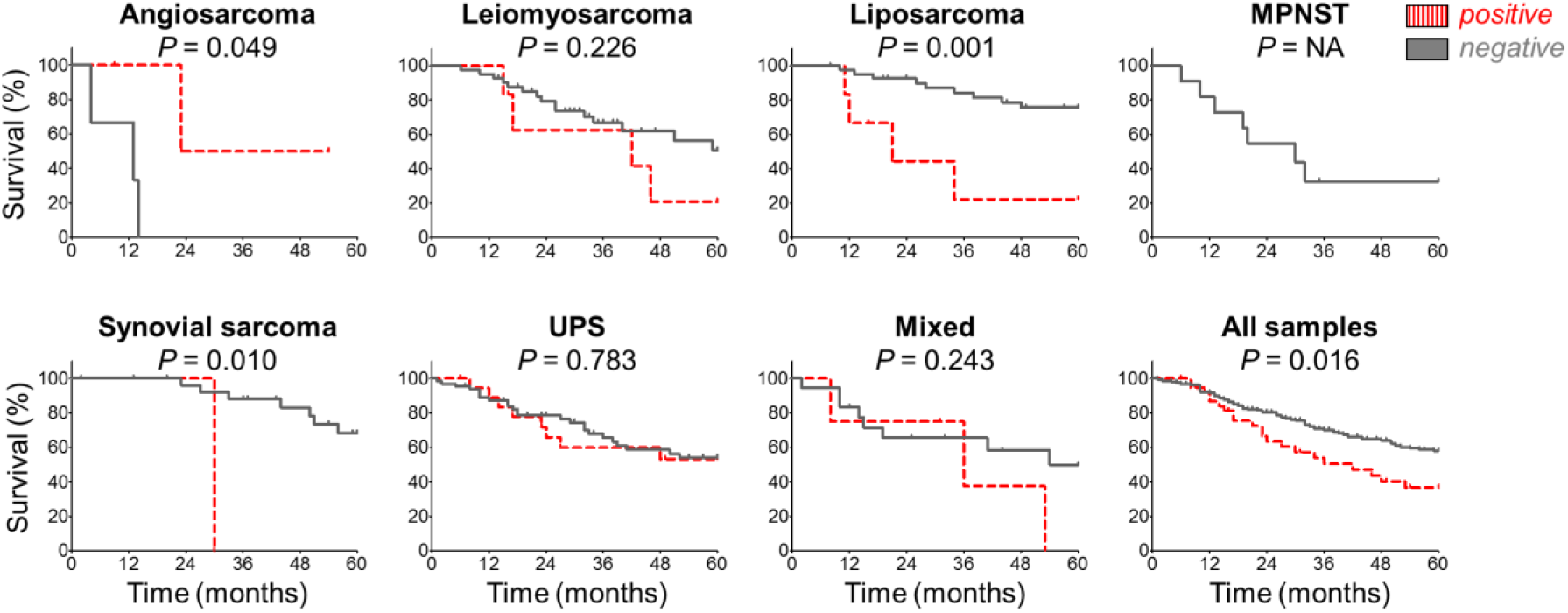
PD-L1 is associated with overall survival in STS. Kaplan-Meier plots indicating the overall survival for the given STS subtypes and for all samples in the entire cohort. Red: PD-L1 positive; grey: PD-L1 negative; significance was assessed by log-rank test.

**Table 4:** Association of PD-L1 expression and clinicopathological parameters in STS

| **STS subtype** | **Sex**<br>**(M/F)** | **Age </ ≥**<br>**median** | **Extr./**<br>**trunk** | **M status**<br>**(M0/M1)** | **Grading**<br>**(1-2/3)** | **Size </ ≥**<br>**80mm** | **Ki-67** | **Survival**<br>**(5y-OS)** |
| --- | --- | --- | --- | --- | --- | --- | --- | --- |
| Angiosarcoma | 1.000<br>(FE) | 1.000<br>(FE) | NA | 1.000<br>(FE) | 1.000<br>(FE) | 1.000<br>(FE) | 0.180<br>(TT) | 0.050<br>(LRT) |
| Leiomyosarcoma | 0.653<br>(FE) | 0.221<br>(FE) | 0.614<br>(FE) | 1.000<br>(FE) | 0.161<br>(FE) | 0.645<br>(FE) | 0.365<br>(TT) | 0.226<br>(LRT) |
| Dediff. Liposarcoma | 0.384<br>(FE) | 0.098<br>(FE) | 0.324<br>(FE) | 0.068<br>(FE) | 1.000<br>(FE) | 0.565<br>(FE) | 0.241<br>(TT) | 0.001<br>(LRT) |
| MPNST | NA | NA | NA | NA | NA | NA | NA | NA |
| Synovial sarcoma | 0.379<br>(FE) | 1.000<br>(FE) | 0.483<br>(FE) | 1.000<br>(FE) | 1.000<br>(FE) | 1.000<br>(FE) | NA | 0.010<br>(LRT) |
| UPS | 0.113<br>(FE) | 0.190<br>(FE) | 0.001<br>(FE) | 0.028<br>(FE) | 0.587<br>(FE) | 0.783<br>(FE) | 0.441<br>(TT) | 0.783<br>(LRT) |
| Mixed | 0.264<br>(FE) | 0.603<br>(FE) | 0.024<br>(FE) | 0.292<br>(FE) | 1.000<br>(FE) | 0.515<br>(FE) | 0.397<br>(TT) | 0.243<br>(LRT) |
| **All samples** | **0.009**<br>**(FE)** | **0.49**<br>**(FE)** | **0.006**<br>**(FE)** | **0.021**<br>**(FE)** | **0.022**<br>**(FE)** | **0.571**<br>**(FE)** | **0.046**<br>**(TT)** | **0.016**<br>**(LRT)** |

FE: Fisher's exact test; TT: Student's t-test; LRT: log-rank test, extr.: extremities; 5y-OS: 5-year overall survival

## DISCUSSION

Immune checkpoint inhibitors interfering with the interaction of tumoral PD-L1 and PD-1 on TILs show promising results as anti-cancer drugs in recent studies, e.g. for melanoma and bladder carcinoma^10,11^. So far, the applicability of these inhibitors for STS patients has not been studied extensively^15^.

Since any targeted therapy is prone to fail in the absence of the target^16^, we first assessed the PD-L1 expression in STS at protein level. To this end, we compiled a cohort of 247 tumor samples from STS patients with matched and well-curated clinical data, including median follow-up of 3 years. The cohort comprised six distinct and representative STS subtypes, including the three most common subtypes in adults (UPS, liposarcoma, leiomyosarcoma) with ≥47 samples, and additional 22 mixed STS cases. Thus, to the best of our knowledge, our analyses were based on the largest and most comprehensive STS cohort to date^13,17^.

For MPNST not any sample was scored as positive for PD-L1, for angiosarcomas 50%, which indicates a strong variability of PD-L1 expression depending on the STS subtype. However, it should be noted that for both subtypes showing extreme variability of PD-L1 only a limited number of samples (11 and 6, respectively) were available.

Interestingly, 23% of UPS cases (19/83) were positive for PD-L1, while only 13% of leiomyosarcomas (6/47) and 12% of dedifferentiated liposarcomas (6/49) showed PD-L1 expression. This is in line with prior studies in other cancers correlating mutational burden, which is much higher in UPS compared to leiomyosarcoma and liposarcoma^18^, with enhanced neoantigen presentation^19,20^. Hence, immune checkpoint inhibition may be in particular effective in UPS.

As PD-L1 acts as ligand of PD-1, we next investigated the positivity for PD-1 expressing TILs in the same patients’ specimens tested for PD-L1 positivity. Across all STS subtypes with ≥6 samples, the rate of PD-1 positive samples in PD-L1 expressing ones was higher than in the negative ones. These results are supported by previous studies in smaller STS cohorts or single STS subtypes demonstrating a PD-L1 and PD-1 interaction^13,14,17,21^, which collectively provide a strong rationale that PD-L1 positive patients are eligible for immune checkpoint inhibitor therapy. Furthermore, our findings support the hypothesis that cancer cells can dynamically increase PD-L1 expression to protect themselves in settings of increased numbers of TILs^12^. Besides the role of PD-L1 as a biomarker to guide clinical decisions on the implementation of immunotherapy, we evaluated its prognostic relevance. In our large STS cohort we found that intratumoral PD-L1 expression was associated with significantly worse 5-year overall survival (*P* = 0.016). Although we intentionally limited our survival analyses to 5-year overall survival to avoid potential biases through death of other causes at later time points after diagnosis, it should be noted that even an unrestricted long-time survival analysis for the entire cohort showed suggestive evidence for an association of PD-L1 expression with worse survival (*P* = 0.074). In fact, PD-L1 was associated with several markers for worse clinical outcome, also with higher rates of proliferation assessed by Ki-67 evaluation. Moreover, PD-L1 was significantly positively associated with G3 grading, which further supports previous findings on superior response to checkpoint inhibitors in tumors with high mutational burden^20^.

The observed prognostic relevance of the PD-L1/PD-1 axis indicates that especially PD-L1 positive STS patients should be considered for treatment with immune checkpoint inhibitors, as especially in these patients long-term tumor control may not be achieved with conventional treatment options.

Collectively, we show for a large and comprehensive STS cohort the abundance of PD-L1, PD-1 and TILs across subtypes and provide evidence for the clinical relevance of PD-L1. We conclude that immune checkpoint inhibitor treatment may constitute a promising approach for a substantial proportion of STS patients that show immunohistochemical evidence for intratumoral PD-L1 expression.

## Abbreviations

HPF –: high power field,
IHC –: immunohistochemistry,
MPNST –: malignant peripheral nerve sheath tumor,
STS –: soft tissue sarcoma,
TIL –: tumor infiltrating lymphocyte,
TMA –: tissue microarray,
UPS –: undifferentiated pleomorphic sarcoma

## Author contributions

MFO analyzed the data, performed statistical analysis, and wrote the paper together with TGPG. EK and TKn scored the IHC staining of PD-L1 and PD-1. FCA, LRP, and FW scored Ki-67 staining. VB, MS, EN, LL, and RI supplied clinical data. AAH carried out statistical analyses. TGPG and TKi provided laboratory infrastructure. TKn conceived the project, drafted the paper, and supervised the analyses together with TGPG.

## ACKNOWLEDGEMENTS & FUNDING

We thank A. Heier, M. Mel and A. Sendelhofert for excellent technical support. The laboratory of TGPG. is supported by grants from the ‘Verein zur Förderung von Wissenschaft und Forschung an der Medizinischen Fakultät der LMU München (WiFoMed)’, by LMU Munich’s Institutional Strategy LMUexcellent within the framework of the German Excellence Initiative, the ‘Mehr LEBEN für krebskranke Kinder – Bettina-Bräu-Stiftung’, the Walter Schulz Foundation, the Wilhelm Sander-Foundation (2016.167.1), the Friedrich-Baur foundation, the Matthias-Lackas foundation, the Dr. Leopold und Carmen Ellinger foundation, the Gert & Susanna Mayer foundation, the Deutsche Forschungsgemeinschaft (DFG 391665916), and by the German Cancer Aid (111886 and 70112257).

